# AI-based search for convergently expanding, advantageous mutations in SARS-CoV-2 by focusing on oligonucleotide frequencies

**DOI:** 10.1101/2022.05.13.491763

**Authors:** Toshimichi Ikemura, Yuki Iwasaki, Kennosuke Wada, Yoshiko Wada, Takashi Abe

**Affiliations:** Nagahama Institute of Bio-Science and Technology, Nagahama, Shiga-ken 526-0829, Japan; Faculty of Engineering, Niigata University, Niigata-ken 950-2181, Japan

**Keywords:** COVID-19, artificial intelligence, zoonotic virus, oligonucleotide, Omicron, Self-Organizing Map, Unsupervised machine learning, SARS-CoV-2, SOM, RNA virus

## Abstract

Among mutations that occur in SARS-CoV-2, efficient identification of mutations advantageous for viral replication and transmission is important to characterize and defeat this rampant virus. Mutations rapidly expanding frequency in a viral population are candidates for advantageous mutations, but neutral mutations hitchhiking with advantageous mutations are also likely to be included. To distinguish these, we focus on mutations that appear to occur independently in different lineages and expand in frequency in a convergent evolutionary manner. Batch-learning SOM (BLSOM) can separate SARS-CoV-2 genome sequences according by lineage from only providing the oligonucleotide composition. Focusing on remarkably expanding 20-mers, each of which is only represented by one copy in the viral genome, allows us to correlate the expanding 20-mers to mutations. Using visualization functions in BLSOM, we can efficiently identify mutations that have expanded remarkably both in the Omicron lineage, which is phylogenetically distinct from other lineages, and in other lineages. Most of these mutations involved changes in amino acids, but there were a few that did not, such as an intergenic mutation.

## Introduction

To confront the global threat of COVID-19 [1-3], a massive number of SARS-CoV-2 genome sequences have been rapidly decoded and promptly released through the GISAID database [4]. To understand this formidable virus from multiple perspectives, we must implement diverse research methods such as artificial intelligence (AI) that can efficiently analyze big data even on a personal computer and support efficient knowledge discovery through intuitive visualization. Unsupervised machine learning can provide new information without relying on particular models or assumptions. We previously established an unsupervised AI, batch-learning self-organizing map (BLSOM), for oligonucleotide composition, which can reveal various new characteristics of a wide variety of genomes by analyzing a large number of genome sequences [5-9].

In viral growth and infection, many host factors (e.g., nucleotide pools, proteins and RNAs) and antiviral mechanisms (e.g., antibodies, cytotoxic T cells and interferons) are involved [10-12]. Since human cells may not present ideal growth conditions for zoonotic viruses that have invaded from nonhuman hosts, efficient growth and transmission should require changes in the viral genome after invasion into the human population. Mutations rapidly expanding in frequency in a viral population are candidates for advantageous mutations. A large number of rapidly expanding mutations have been particularly concentrated in the S gene encoding the S glycoprotein, which plays essential roles in viral attachment, fusion and entry into host cells; additionally, its surface location renders it a direct target for host immune responses, making it the main target of neutralizing antibodies [13]. Of course, there are mutations outside the S gene that expand rapidly, including those that do not involve amino acid changes [2,3,14].

Advantageous mutations without amino acid changes have been discussed from multiple perspectives, including interactions with host factors such as RNA-binding proteins [15] and miRNAs [16,17], as well as the higher-order structure of viral RNA [18]. It is thus important to compile examples of remarkably expanded mutations, even outside the S gene, for the development of antiviral drugs, e.g., oligonucleotide drugs. By analyzing oligonucleotide composition to study the viral adaptation, we previously found time-series directional changes (i.e., monotonic increases or decreases) in mono- and oligonucleotide composition of four zoonotic RNA viruses (influenza virus [19-22], Zaire ebolavirus [21,22], MERS coronavirus [21] and SARS-CoV-2 [23-25], which were detectable even on a monthly basis.

In the present study, we used a similar approach to analyze the recently prevalent Omicron subvariants and compare them with previously prevalent lineages. The Omicron lineage is derived from the G clade, but the accumulated mutations are very different from those in other G-derived lineages, indicating that separation from the other clades occurred fairly early in the COVID-19 pandemic [26,27]; it has even been predicted that a certain G-derived virus once spread to a nonhuman animal and then reinvaded the human population [28-30]. If a mutation that occurs independently in different lineages remarkably increases its population frequency commonly in distantly related lineages, convergent evolutionary increase is highly likely due to the increase in fitness caused by the mutation, and therefore, Omicron is believed to be especially well suited for searching for and studying convergently expanding, advantageous mutations that have rapidly spread in the human population [31].

We previously analyzed time-series changes in composition of short and long oligonucleotides from SARS-CoV-2 strains isolated mainly in the first year of the pandemic and found many long oligonucleotides (e.g., 20-mers) that had expanded remarkably in the viral population, which allowed us to predict candidates of advantageous mutations that had spread among humans [9,23,24]. In the present study, we analyzed 20-mer frequencies to search for mutations that expanded markedly in the Omicron population.

## Materials and methods

### Human SARS-CoV-2 genome sequences

SARS-CoV-2 genome sequences of 12 G-derived lineages other than Omicron (Alpha, Beta, Gamma, Delta, Epsilon, Zeta, Eta, Theta, Iota, Kappa, Lambda and Mu) and of 3 other non-G lineages (L, S and V clades) were obtained from the GISAID database (https://www.gisaid.org/epiflu-applications/next-hcov-19-app/) [4]; sequences that were complete, high coverage and from humans were downloaded on 29 November 2021.

For each lineage, the number of isolates per month was counted, and only the genomes with the highest monthly numbers were used for a BLSOM analysis after removing their poly(A)-tails. It should be pointed out that the differences in numbers of viral strains among lineages is extremely large. This makes it difficult to properly understand the characteristics of each lineage in the BLSOM analysis, so for lineages with over 10000 genomes, 10000 were randomly selected.

In the case of Omicron, the pandemic is still ongoing during the course of this study and preparation of this paper, and a large number of genome sequences have been released by GISAID. This appears to make research difficult but offers a distinct advantage in that the results obtained at an early stage of the study can be verified in the same publication with new sequences obtained later, increasing the reliability of the analysis. We adopted this strategy in the case of Omicron. On 19 January 2022, we first downloaded the complete and high-coverage category sequences by month of virus isolation, as done for other lineages. For the isolates for Dec. 2021 and Jan. 2022, 3191 and 3701 sequences were obtained, respectively, while for Nov. 2021, only 17 sequences were obtained most likely due to difficulties in sequencing of genomes that were very different from known sequences in the early stages. Information at the beginning of the Omicron pandemic was thought to be important, but 17 sequences were too few to adequately know their features. For the low-coverage category, the 2299 sequences, which included BA.3 sequences, were obtained. Thus, in the BLSOM analysis, which was intended to provide an overall picture of the Omicron lineage, the 2299 sequence was used for the Nov. 2021 isolates. The sequences downloaded on 29 January 2022 were used to verify the degree of separation between BA.1 and BA.2 on BLSOM, and finally, the sequences downloaded on 3 March 2022 were used for verification of the peculiarity of the 1169 20-mers of interest in the Omicron lineage.

### BLSOM

The self-organizing map (SOM) developed by Kohonen [32,33] is an unsupervised neural network algorithm that implements characteristic nonlinear projection from the high-dimensional space of input data onto a two-dimensional array of weight vectors. Kanaya et al. [5] modified the conventional SOM for genome informatics on the basis of batch learning (BLSOM), aiming to make the learning process and the resulting map independent of the order of data input. In the original SOM, the initial vectorial data were set by random values, but in the BLSOM, the initial vectors were set based on the widest scale of sequence distribution in the oligonucleotide frequency space with PCA (principal component analysis). In the oligonucleotide BLSOM in the present study, the number of nodes (lattice points) was set to 1% of the total number of sequences used. The weight vectors (wij) were arranged in a two-dimensional lattice denoted by i (= 0, 1,…, I-1) and j (= 0, 1,…, J-1). Weights in the first dimension (I) were arranged into lattices corresponding to a width of five times the standard deviation (5σ1) of the first principal component, and the second dimension (J) was defined by the nearest integer greater than σ2/σ1 × I; σ1 and σ2 were the standard deviations of the first and second principal components, respectively. Weight vectors were updated as described previously [7]. A BLSOM program is available on our website (http://bioinfo.ie.niigata-u.ac.jp/?BLSOM).

## Results and discussion

### Histogram analysis of markedly expanded 20-mers in the Omicron lineage

Omicron variants first appeared in Nov. 2021 and have since caused a global pandemic [26-29]. In the early stages of the Omicron pandemic in 2021, BA.1 sublineage was prevalent, but the BA.2 sublineage began to dominate worldwide in 2022 [34,35] and (https://covariants.org/ [36]). In this study, we first analyzed the Omicron genome sequences downloaded on 19 January 2022. To obtain up-to-date information on both BA.1 and BA.2, we first focused on 3701 strains that were isolated in Jan. 2022 and belonged to the complete and high-coverage category in GISAID; for details, refer to Materials and methods. In the viral genome, there was one copy of each 20-mer excluding the poly(A)-tail, which was removed before the analysis. Thus, analysis of 20-mers that remarkably increased in frequency in the Omicron population after onset of the SARS-CoV-2 pandemic can provide information on mutations whose population frequencies have increased markedly.

After calculating the frequency of each 20-mer in the Jan. 2022 population (3701 strains), we calculated the difference from the corresponding frequency in the viral population in the first month of the pandemic, which was presented by 22 strains isolated in Dec. 2019. Then, we focused on 20-mers that had increased in frequency in the Jan. 2022 population compared to the Dec. 2019 population. The level of increase was calculated for each 20-mer, and the 20-mers were divided into 0.05 increments of proportion by increase level; the number of 20-mers in each division is display as a histogram in Fig 1A.

**Fig 1.**
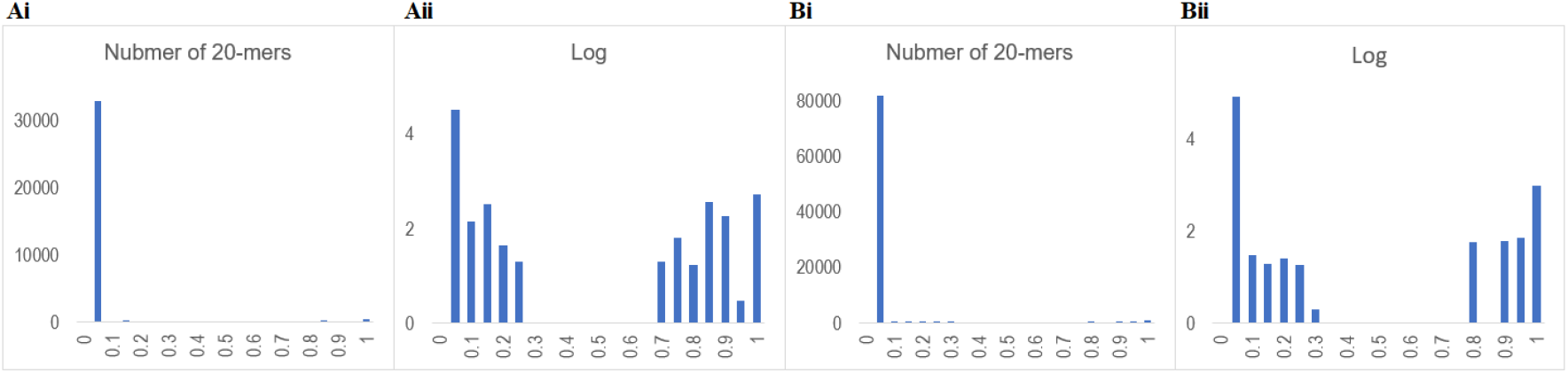
Histogram analysis of 20-mers. (A) Histogram of the level of increase of each 20-mer frequency in the Jan. 2022 population compared to that in the Dec. 19 population. The horizontal axis is divided by the increase level by 0.05. The vertical axis shows normal numbers (i) or logarithms (Log) (ii). Here, nonexistence in the logarithmic display is shown expediently as 0. (B) The level of increase of each 20-mer frequency in the Feb. 2022 population compared to that in the Dec. 19 population is displayed as described in A.

In Fig. Ai, the horizontal axis is divided by 0.05 increments in proportional representation of each level of increase, and the vertical axis shows the number of 20-mers belonging to each division. The highest peak near 0 on the horizontal axis show that most of new 20-mers generated by mutations have not significantly expanded in frequency. However, a very low vertical bar can be seen near the horizontal axis of 1, and the logarithmic display is shown in Fig 1Aii. Distinct peaks of 20-mers are observed in the high abscissa region; the 20-mers in the peaks near 1 correspond to those present in most strains in the Jan. 2022 population. Interestingly, there is no peak between 0.3 and 0.65 on the horizontal axis, showing that the distribution is completely separated. It should be noted that in this display log 0 is set to 0 for convenience. Since there is no division having a value of 1, there is no confusion, and there are no 20-mers between 0.3 and 0.65.

The 20-mers in the higher-frequency side in Fig 1Aii, which are the 1169 20-mers held in 67% or more of the Omicron strains, are considered to be the remarkably expanded 20-mers in the Omicron lineage. Since these 20-mers were absent at the beginning of the epidemic, they emerged by mutations and spread markedly. These are noted “1169 Omicron 20-mers” and used in the subsequent analysis. The results presented in Fig 1B will be explained in the last part of this paper in connection with the distinctive features of the 1169 20-mers in the Omicron lineage.

### BLSOM analysis with 1169 Omicron 20-mers

We next performed a BLSOM analysis to examine whether the 1169 Omicron 20-mers also expanded in other 15 lineages and, if so, to what extent; details of genome sequences used for the 15 lineages are explained in Materials and methods. For each genome in each of 16 lineages, including Omicron, we first calculated the frequency of occurrence of the 1169 Omicron 20-mers and constructed a BLSOM using the frequency per genome as vectorial data (Fig 2A). The number of genomes for each lineage used in BLSOM is given in the Fig 2 legend. The total number of nodes (i.e., grid points) was set to approximately 1/100 of the total number of viral genomes (81,229). Importantly, no information other than the 20-mer occurrence frequency was provided during the machine learning step: unsupervised AI. After the machine learning, to determine whether the clustering (self-organization) achieved by the BLSOM corresponded to the known lineages, nodes containing genomes of a single lineage were colored to indicate each lineage, and those containing genomes of multiple lineages were displayed in black. It should be noted that when nodes that have sequences with distinctly different frequencies are located nearby, empty nodes, to which no sequences are attributed after the learning, often appear between them, and these nodes are left blank (no color) on the BLSOM [8,9]. As in our previous BLSOM analysis of SARS-CoV-2 strains [9,23,24], which were mainly isolated within the first year of the pandemic, good clustering by lineage was obtained, and Omicron strains formed their own territory (red) on the right side of the map (Fig 2A).

**Fig 2.**
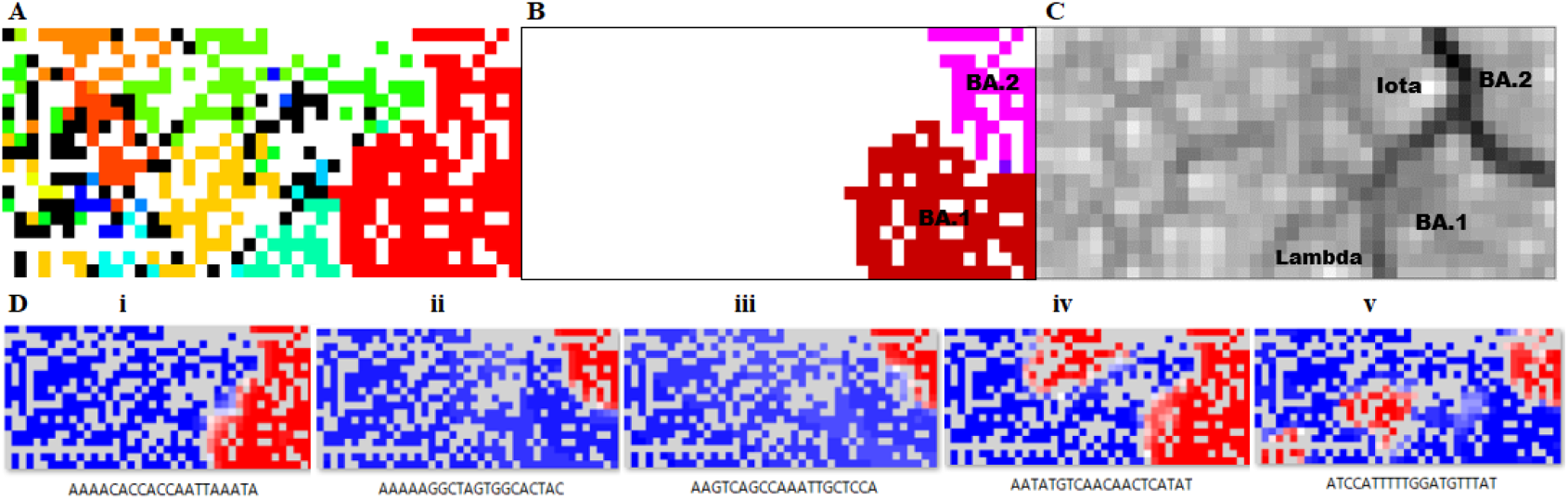
BLSOM for 1169 Omicron 20-mers. (A) The total number of nodes (grid points) was set to approximately 1/100 of the total number of genomes: 817 nodes. Nodes that included sequences from more than one lineage are indicated in black, and those containing sequences from a single lineage are shown in the following lineage-specific color with the number of genomes in parentheses: Omicron 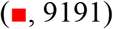, Alpha 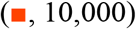, Beta 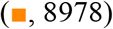, Delta 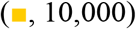, Epsilon 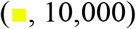, Eta 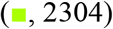, Gamma 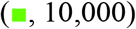, Iota 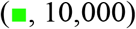, Kappa 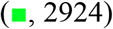, L 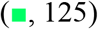, Lambda 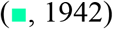, Mu 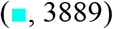, S 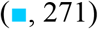, Theta 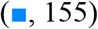, V 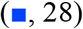 and Zeta 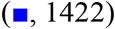. Nodes that included no sequences were left in blank (white). (B) Mapping of newly downloaded BA.1 and BA.2 sequences on the BLSOM presented in A. (C) U-matrix. BA.1, BA.2, Alpha and Iota territories, which are visually separated by black lines of the U-matrix, are specified. (D) Heatmaps for five 20-mers. Those for all 1169 20-mers are presented in S1 Fig.

As mentioned above, the subdivision of Omicron strains into BA.1 and BA.2, the latter of which has become prevalent in 2022, and BA.3, which caused a small outbreak in 2021, is noted in GISAID; for sequences of the sublineage strains, refer to Materials and methods. The BLSOM in Fig 2A included all Omicron strains isolated from Nov. 2021 to Jan. 2022, so we can examine territories of these three sublineages on the BLSOM by mapping the strains in the three sublineages separately onto the BLSOM (Fig 2B). In more detail, using the vectorial data of the occurrence frequency in each genome, we searched for the node with the closest Euclidean distance, attributed each genome to that node and colored the node for each sublineage, as described previously [7,9]. Separation between BA.1 (reddish brown) and BA.2 (pink) was clear, and BA.3 (violet), which remained in a small population, was in the middle of the former two sublineages (Fig 2B). It is also clear that no Omicron strain was located in any other lineage territories. The Euclidean distance between representative vectors of neighboring nodes can be visualized by the degree of blackness with the U-matrix [37]. Fig 2C shows the U-matrix pattern of the BLSOM in Fig 2A, with a greater Euclidean distance reflected by a higher degree of blackness. Notably, the boundary between Omicron and other lineages and that between BA.1 and BA.2 are represented by dark black lines (Fig 2C), showing that the oligonucleotide frequencies are very different between these; territories of BA.1 and BA.2 are illustrated in the U-matrix. Two G-derived groups, Lambda and Iota, are adjacent to the Omicron territory, and these are also illustrated.

### 20-mers that contributed to clustering by lineage

BLSOM can explain diagnostic oligonucleotides responsible for lineage-dependent clustering with the following heatmaps. The representative vector for each node of the BLSOM presented in Fig 2A is composed of 1169 variables, and the contribution level of each variable at each node can be visualized by the heatmap [6]: high (red), moderate (white) and low (blue). In this red/blue heatmap, empty nodes, which had no sequence in the BLSOM, are indicated in gray. All 1169 heatmaps are presented in S1 Fig along with 20-mer sequences, and typical examples are shown in Fig 2D. Fig 2Di shows an example of heatmaps where only the entire region of the Omicron territory is red: a 20-mer retained in almost all Omicron genomes. Fig 2Dii shows a heatmap where only the entire region of BA.2 is red: a 20-mer retained in almost all BA.2 genomes. Fig 2Diii shows a heatmap where only a part of BA.2 is red: a 20-mer retained in a portion of BA.2 genomes. Fig 2Div and v shows an example of 20-mers retained by the majority of Omicron and BA.2 plus non-Omicron lineages, respectively, which are candidates for mutations that have increased convergently in both Omicron and non-Omicron lineages and thus form the main focus of this study. Notably, as is clear from Supplementary S1 Fig, there are a wide variety of patterns, and efficient data mining from many images with various patterns requires a systematic approach. We next introduce an AI method to search for patterns of interest and classify them.

### Clustering of heatmap patterns by BLSOM

AI is widely used for image classification, and we previously developed a BLSOM for clustering a large number of heatmaps derived from the oligonucleotide BLSOM [9]. Here, we applied the BLSOM to 1169 heatmaps presented in Supplementary S1 Fig. As a typical instance of AI image processing, the 2D heatmap pattern for each 20-mer was converted to one-dimensional vector data; the number of variables in each vector is the number of nodes containing genome sequences in the BLSOM illustrated in Fig 2A. In more details, we focus on 471 variables, each of which represents the contribution level of each 20-mer in the node.

In this BLSOM for heatmaps, the number of nodes was set to 24 (approximately 50 heatmap patterns per node). Here, nodes having multiple 20-mer patterns are shown in black, while nodes with no pattern are left blank (Fig 3A). The number of patterns that were attributed to each node is shown by the height of a colored column in a 3D display (Fig 3B), and for each column, a representative example of heatmaps attributed to the corresponding node is presented. First, the green column on the rightmost will be described. The heatmap attributed to this column is red for all G-derived lineages including Omicron, but blue for all non-G clades (L, S and V), showing that the original G strain, from which Omicron was derived, already had the corresponding 20-mers. It should be noted here that eighty similar heatmaps belonged to this column, and the corresponding eighty 20-mers could be classified into four groups of 20-mers as described later.

**Fig 3.**
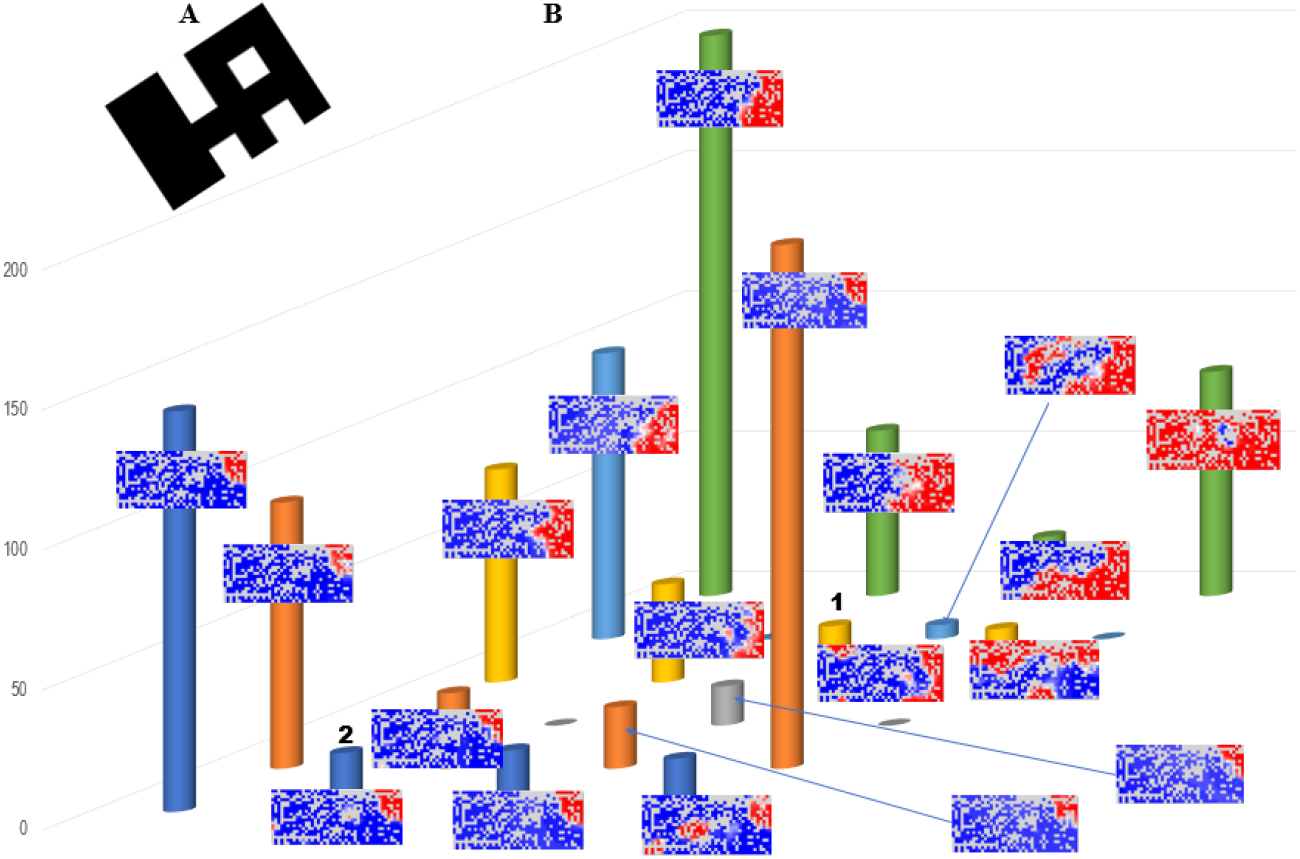
BLSOM for heatmap patterns. (A) BLSOM of 1169 heatmaps from the BLSOM presented in Fig 2A. Nodes including more than one heatmap are indicated in black, and those that include no patterns were left in blank (white). The 2D patten is deliberately tilted to show the relationship with the following 3D display in an easy-to-understand manner. (B) 3D display of the BLSOM presented in A. The number of patterns belonging to each node is indicated by the height of the colored column, and for each column, a representative example of heatmaps attributed to the corresponding node is shown.

We next describe the highest green column, which is at the left end of the far side and consists of approximately 200 heatmaps. In these heatmaps, the whole area of Omicron (BA.1+BA.2) is red, and thus, the highest column is composed of the 20-mers present in almost all Omicron strains; columns with similar patterns are located near it. The second highest, brown column locating on the near side is composed of 20-mers present in almost all BA.2 strains. As mentioned when explaining the histogram analysis illustrated in Fig 1, we are focusing on 1169 different 20-mers with frequencies at least 67% in the Jan. 2022 population, and the 20-mers with relatively low frequencies show various patterns of absence in the BA.2 territory and form multiple columns located mainly on the near side.

In this study, we are particularly interested in patterns in which regions other than the Omicron territory also show red color, such as those presented in Fig 2Div and v.

Patterns in which red color is observed in the BA.1 and non-Omicron territories are located at the front, and those in which red color is observed in the BA.1+BA.2 and non-Omicron territories are located at the back. The heights of the columns with these patterns were relatively low, with most having approximately 20 patterns or integer multiples of that. Importantly, 20-mers attributed to each column were found to consist of sequences that were shifted by one base, as described below.

### Association of focal 20-mers with mutations

It should be mentioned again that 1169 Omicron 20-mers were not present at the epidemic start (Dec. 2019) and thus were caused by mutations. To relate the focal 20-mers to mutations, we first selected one genome sequence each from BA.1 and BA.2 that was particularly long and had few unknown nucleotides (Ns): GISAID Accession numbers EPI_ISL_7336152 for BA.1 and EPI_ISL_8465178 for BA.2. Using these sequences as a template, we performed a BLASTn search of the 20-mers that belonged to columns with red regions in territories of both Omicron and other lineages. We first explain the results obtained by taking as an example the eighty 20-mers belonging to the green column on the rightmost side in Fig 3B. The BLASTn search divided the eighty 20-mers into four groups, with each group consisting of twenty 20-mers that were shifted by one base. In other words, these eighty 20-mers were found to be components of four 39-mers. In the case of long sequences such as 39-mers, a BLASTn search against a standard genome sequence of a SARS-CoV-2 strain isolated in Dec. 2019 (NC_045512 in GenBank) [38,39] allowed alignment even when there were multiple mutations in close proximity including indels; the presence of these complex mutations yielded the assembled sequence other than the 39-mer. The mutations found in this BLASTn search against the standard sequence were consistent with known mutations in Omicron (https://covariants.org/) [36]. In the case of the eighty 20-mers attributed to the rightmost green column, all four 39-mers had a single nucleotide mutation in the center, and these were the diagnostic mutations that characterized all G lineages, including Omicron: C241T in 5’ UTR, C3037T and C14408T in ORF1ab and A23403G in the S gene [14, 23]. However, the main focus of this study is on mutations that occurred after Omicron and other G-derived lineages diverged, i.e., that Omicron shares with some of other lineages.

### Mutations shared between Omicron and some of other lineages

Table 1 lists the mutations that Omicron shares with some of other lineages along with the territories in which the mutated 20-mers were located in the BLSOM in Fig 2A. To distinguish qualitatively the range occupied by red in each territory, we used +++ to indicate that red nodes were found in almost the entire area of a certain lineage territory as for Fig 2Di, the related mutations are marked with +++ in both BA.1 and BA.2. In a case where a portion of the red region is clearly missing in a particular territory as for the example numbered 1 in Fig 3B, the corresponding mutation is marked with ++ for the territory. If the red region was extremely limited as for the example numbered 2 in Fig 3B, the corresponding mutation is specified with +; even for the + category, a few or several hundreds of genome sequences were usually found because the condition of BLSOM (Fig 2A) was set so that an average of 100 sequences belonged to each node, indicating that the corresponding mutations expanded to an unignorable level in the corresponding lineages.

**Table 1.** Mutations shared between Omicron and some of other lineages.

| Mutation | Gene | Product | A.A change | BA.2 | BA.1 | Lambda | Iota | Delta | Mu | Alpha | Beta | Gamma | Epsilon | S | L |
| --- | --- | --- | --- | --- | --- | --- | --- | --- | --- | --- | --- | --- | --- | --- | --- |
| C10029T | ORF1ab | nsp4 | T492I | +++ | +++ | +++ |  | ++ | ++ |  |  |  |  |  |  |
| 11288_11296del | ORF1ab | nsp6 | SGF3675del | +++ |  | +++ | +++ |  |  | +++ |  |  |  |  |  |
| C23604A | S | spike | P681H | +++ | +++ |  |  |  | ++ | ++ | + |  | + |  |  |
| C28311T | N | nucleocapsid | P13L | +++ | +++ | +++ | ++ |  |  |  |  |  |  | + |  |
| G22813T | S | spike | K417N | +++ | ++ |  |  | + | + |  | ++ |  |  |  |  |
| G22992A | S | spike | S477N | +++ | ++ |  | ++ |  |  |  | + | + |  |  |  |
| A28271T | Intergenic* |  |  | +++ | +++ | +++ |  |  |  |  |  |  |  |  |  |
| C23525T | S | spike | H655Y | +++ | +++ |  |  |  |  |  |  | ++ |  |  |  |
| C22995A | S | spike | T478K | +++ | ++ |  |  | ++ |  |  |  |  |  |  |  |
| A23013C | S | spike | E484A | +++ | ++ |  |  |  |  |  |  |  |  |  |  |
| A23055G | S | spike | Q498R | +++ | ++ |  |  |  |  |  |  |  |  |  |  |
| A23063T | S | spike | N501Y | +++ | ++ |  |  |  |  |  |  |  |  |  |  |
| 28363_28374del | N | nucleocapsid | ERS31del | +++ | ++ |  |  |  |  |  |  |  |  |  |  |
| C17410T | ORF1ab | nsp13 | R5716C | +++ |  |  |  |  |  |  |  |  | + |  | + |
| C9344T | ORF1ab | nsp4 | L3027F | +++ |  |  |  |  |  |  | + |  |  |  |  |
| C12880T | ORF1ab | nsp9 |  | +++ |  |  | + |  |  |  |  |  |  |  |  |
| C21618T | S | spike | T19I | +++ |  |  |  | + |  |  |  |  |  |  |  |
| C22792T | S | spike |  | +++ |  |  |  | + |  |  |  |  |  |  |  |
| C26060T | ORF3a | ORF3a | T223V | +++ |  |  |  | + |  |  |  |  |  |  |  |

\*Intergenic between ORF8 and N

The lineages in Table 1 are ordered in the order of the sum of the qualitative indicators of + for each lineage. BA.2 appeared first because it had all mutations that Omicron shared with others. Concerning lineages other than Omicron, Lambda, which was first isolated in Peru, was highest and followed by Iota, which was first isolated in North America. The order of 19 mutations was in the order of the sum of + for each mutation. Since 16 of the 19 mutations involved changes in amino acids, the mutations thus selected are considered to affect viral functions. Concerning the other three, A28271T is a mutation that does not cause an amino acid change and is located just before the N gene. Importantly, this intergenic mutation is found in almost all strains of the Lambda lineage (+++), indicating that the mutation may have an advantageous effect possibly by regulating the production level of the N protein. The other two (C12880T and C22792T) are synonymous mutations, C to T, which occur frequently due to APOBEC5 enzymes [40,41]. Of the vast number of known C to T mutations, these two remained in the present analysis, making them candidates for advantageous mutations, albeit with only slight advantage. Although + symbols and their totals are qualitative indicators, these appear to provide a certain indication of advantageousness.

The following point should be noted here. When we focus on a certain remarkably expanding mutation, there are no 20-mers representative only of the focal mutation if other remarkably expanding mutations occur within 10 bases on its either side, but only 20-mers representative of the pair with the neighboring mutation. Although there may be examples where it becomes important to treat neighboring mutations as a combination [42], in general it is appropriate to treat them independently. Shortening 20-mer to 10-mer or even less entails the complexity of multiple copies in the genome, but to some extent this problem would be avoided [23]. However, there are obvious limitations for shortening the oligonucleotide. The present method is suitable for excavating advantageous mutations in regions that have received less attention, rather than in the S gene region where the concentration of functionally influential mutations is apparent.

Since the mutations in Table 1 cover rather randomly a wide range of lineages, most (if not all) of the mutations are thought to have expanded in a convergent evolutionary way. We will here discuss a reason that Lambda and Iota were more closely related to Omicron than other lineages, both in the BLSOM and in Table 1. Mutations presented in Table 1 are thought to have occurred far prior to the time when the Omicron genome sequence was first reported (Nov. 2021), because almost all Omicron strains isolated in the Nov. 2021 had these mutations. In the case of coronaviruses, recombination plays an important role in their evolution, and therefore, it is difficult to identify the true origin of each mutation in focus. The proximity of Omicron to Alpha and Iota in the BLSOM and in Table 1 may not reflect their evolutionary origins.

### Analysis of the newly accumulated Omicron genome sequences

One peculiarity of SARS-CoV-2 research is that, regardless of when genome sequences were obtained for analysis, a large number of new sequences have accumulated during the analysis and manuscript preparation; in the present study, the accumulation of genome sequences of Omicron, especially the BA.2 sublineage, was remarkable. This peculiarity appears to make the research difficult but does provide a distinct advantage in that the results obtained at an early stage of study can be verified with new sequences obtained later in the same study, increasing the reliability of the analysis: a cycle of prediction and verification in one evolutionary study. In the final portion of this paper, we took advantage of this situation and examined the peculiarity of the 1169 Omicron 20-mers on which we have focused. In Fig 1B, a histogram analysis was conducted using 32,482 strains isolated in Feb. 2022, which were ten times more than and totally different from those used in Fig 1A. Distinct divisions were again seen; there were no 20-mers between 0.25 and 0.71. Importantly, the number of 20-mers above this 0.71 was 1169, as before, and their sequences were perfectly matched, confirming the peculiarity of the 1169 Omicron 20-mers in the Omicron lineage.

Next, we explain the contents of the highest peak near abscissa 0 in Fig 1Bi. It consists of more than 235,000 20-mers, but over half of them have appeared only in 1∼2 out of 32,482 strains, and none of them were present at the onset of the pandemic. Although a minor portion might be caused by misassignments for sequencing, most should be caused by mutations. Since approximately 90% of 20-mers appear in only 10 strains or less, a major portion of mutations have not spread significantly within the viral population. If one mutation is roughly assumed to lead to twenty 20-mers, more than 11,000 mutations will be found in the over 32,000 genomes analyzed here, and most of them are considered functionally neutral. Importantly, only a very small portion of mutations were remarkably expanding within the Omicron population, and the expanding set, the 1169 Omicron 20-mers, was the same, even analyzing totally independent sequences. This proves their evident peculiarity in the Omicron lineage.

In the study of viral evolution, the use of phylogenetic tree construction methods is undoubtedly the mainstream of research, but it is also important to develop a variety of other methods, including AI, to combat the threat of infectious viruses. The BLSOM introduced here is capable of analyzing large amounts of genome sequences on a personal computer, enabling efficient knowledge discovery that is easy to understand visually. To further promote the study of viral evolution as an interdisciplinary issue, we believe that the active introduction of various AI technologies is important.

## Supporting information

**S1 Fig. Heatmap patterns**. All 1169 heatmaps are presented along with 20-mer sequences.

## Acknowledgments

We gratefully acknowledge the authors submitting their sequences from GISAID’s Database. This work was supported by JSPS KAKENHI Grant Number 18K07151 and JST CREST Grant Number JPMJCR20H1.

## Author Contributions

T. Ikemura and T. Abe designed this project and Y. Iwasaki conducted genome analysis. K. Wada, Y. Wada and Y. Iwasaki developed computer programs. T. Ikemura wrote the manuscript. All authors participate in the discussion through the project.

