## Supplementary figures and images for "AI-based search for convergently expanding, advantageous mutations in SARS-CoV-2 by focusing on oligonucleotide frequencies"

### S1 Fig

**S1 Fig. Heatmap patterns.** All 1169 heatmaps are presented along with 20-mer sequences.

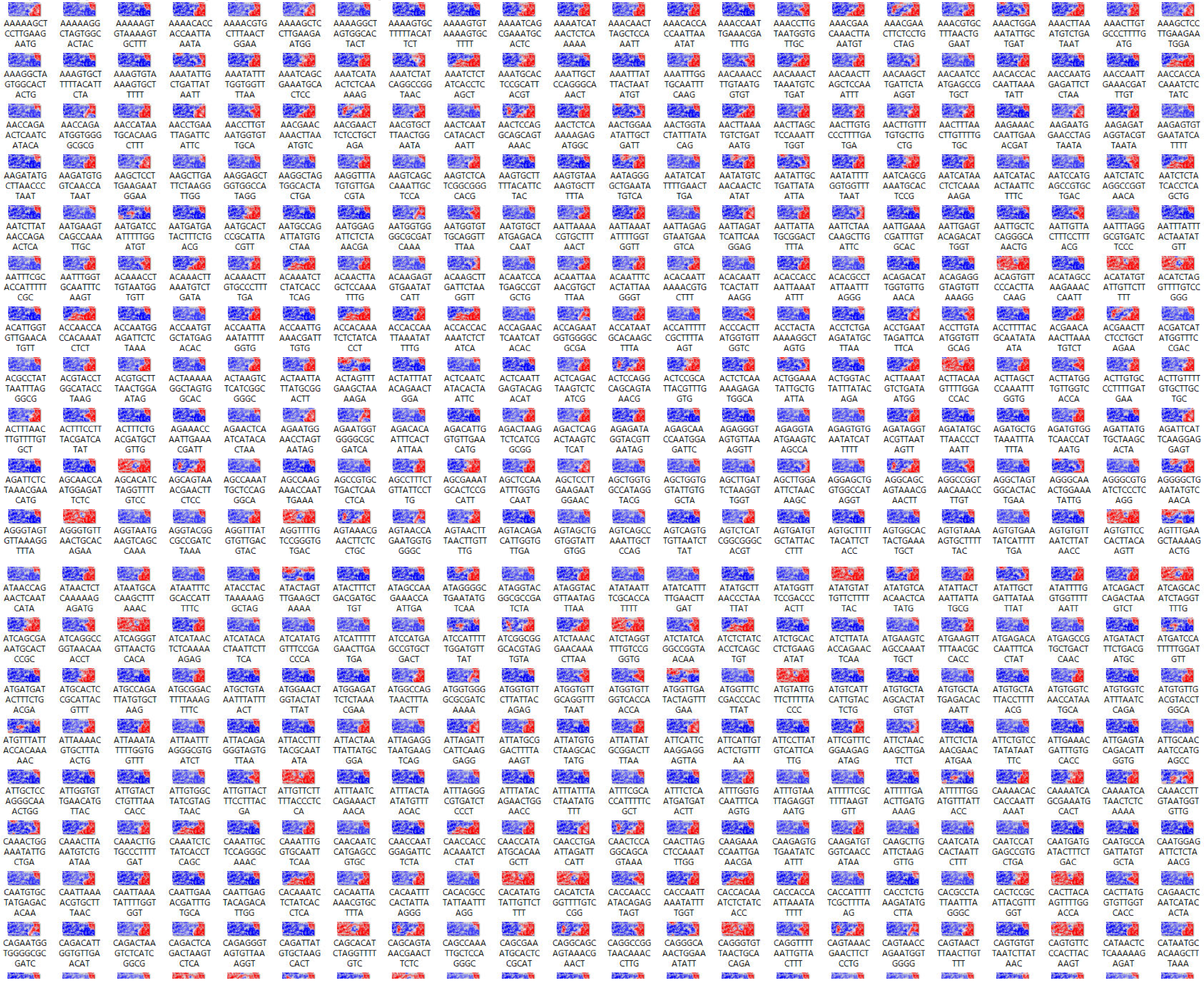

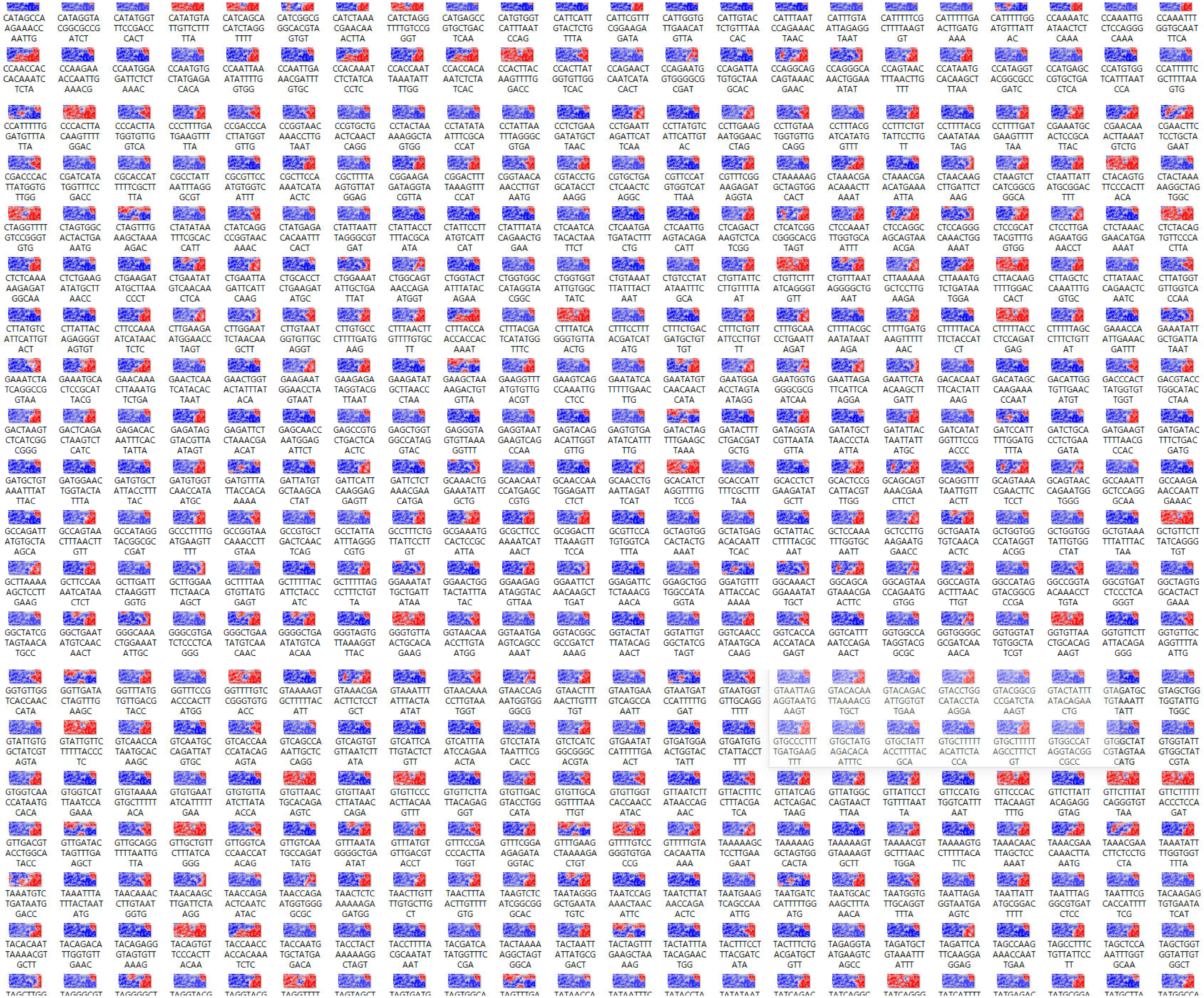

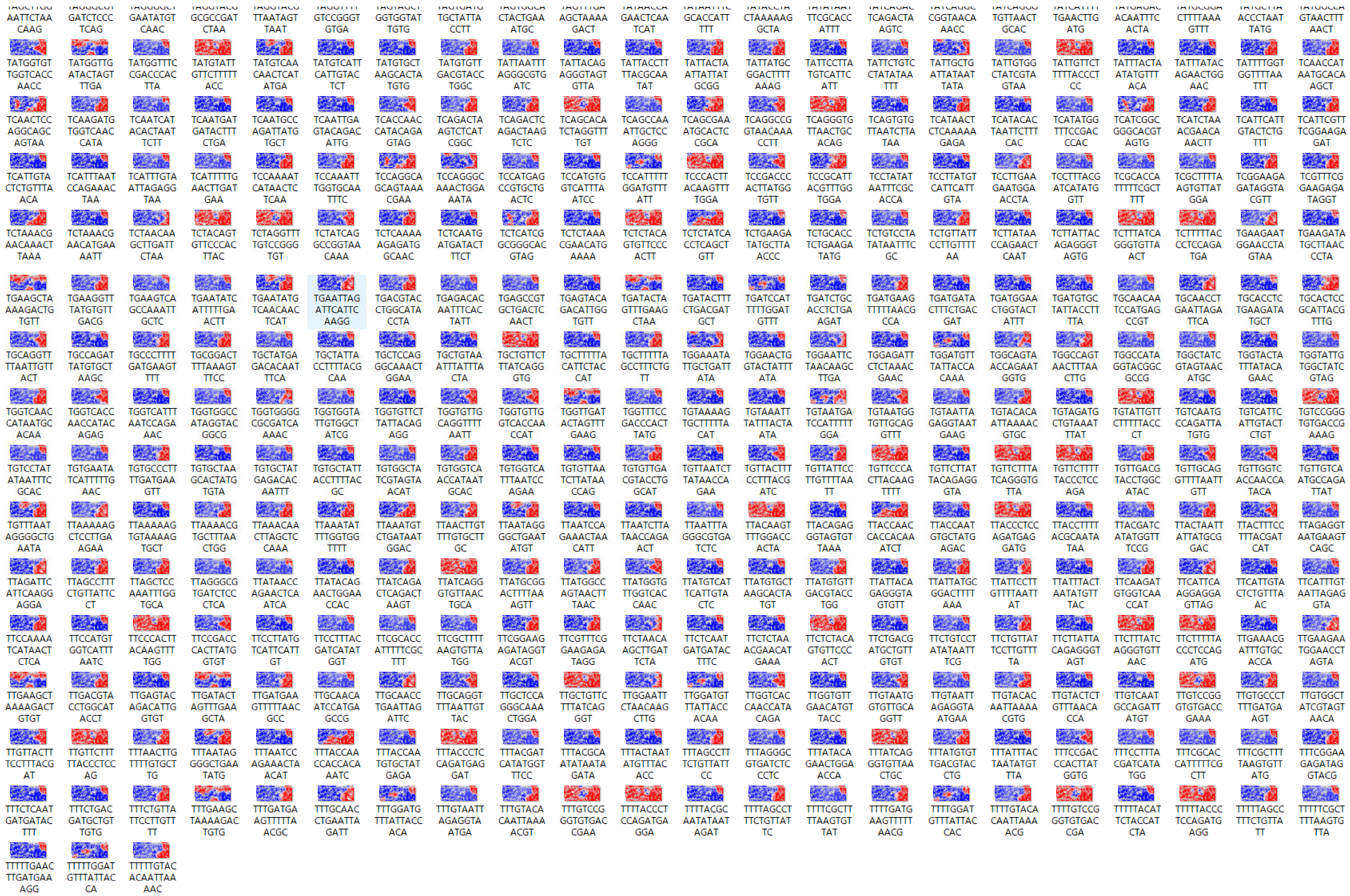
